# Stress Knowledge Map: A knowledge graph resource for systems biology analysis of plant stress responses

**DOI:** 10.1101/2023.11.28.568332

**Authors:** Carissa Bleker, Živa Ramšak, Andras Bittner, Vid Podpečan, Maja Zagorščak, Bernhard Wurzinger, Špela Baebler, Marko Petek, Maja Križnik, Annelotte van Dieren, Juliane Gruber, Leila Afjehi-Sadat, Anže Županič, Markus Teige, Ute C. Vothknecht, Kristina Gruden

**Affiliations:** Department of Biotechnology and Systems Biology, National Institute of Biology, Večna pot 121, SI-1000 Ljubljana, Slovenia; Plant Cell Biology, Institute of Cellular and Molecular Botany, University of Bonn, Kirschallee 1, D-53115 Bonn, Germany; Department of Knowledge Technologies, Jožef Stefan Institute, Jamova cesta 39, SI-1000 Ljubljana, Slovenia; Department of Functional & Evolutionary Ecology, University of Vienna, Djerassiplatz 1, AT-1030 Vienna, Austria; Mass spectrometry unit, Core Facility Shared Services, University of Vienna, Djerassiplatz 1, AT-1030 Vienna, Austria

**Keywords:** knowledge graph, database, plant stress responses, plant signalling, systems biology, digital plant

## Abstract

Stress Knowledge Map (SKM, https://skm.nib.si) is a publicly available resource containing two complementary knowledge graphs describing current knowledge of biochemical, signalling, and regulatory molecular interactions in plants: a highly curated model of plant stress signalling (PSS, 543 reactions) and a large comprehensive knowledge network (CKN, 488,390 interactions). Both were constructed by domain experts through systematic curation of diverse literature and database resources. SKM provides a single entrypoint for plant stress response investigations and the related growth tradeoffs. SKM provides interactive exploration of current knowledge. PSS is also formulated as qualitative and quantitative models for systems biology, and thus represents a starting point of a plant digital twin. Here, we describe the features of SKM and show, through two case studies, how it can be used for complex analyses, including systematic hypothesis generation, design of validation experiments, or to gain new insights into experimental observations in plant biology.

## Introduction

The already apparent effects of climate change on agriculture (Shukla *et al*.), the spread of pests into new regions (Garrett, 2013; IPPC Secretariat, 2021), and rapid population growth (UN DESA, 2022) provide immediate challenges to global food security (Steinwand and Ronald, 2020). Projections show that in order to meet 2050 demand, an increase in crop production of up to 75% is required (Hunter *et al*., 2017). This can be achieved with yield improvements through the development of stress resilient crops, a process requiring a holistic understanding of the effect of stressors on plants. The rapid development of modern ‘omics’ technologies allows for the generation of large and complex datasets, characterising system wide responses. To understand the biological meaning of these large-scale data sets and generate meaningful hypotheses, contextualisation within current knowledge is needed. We have assembled an integrated resource of plant signalling, **S**tress **K**nowledge **M**ap (SKM, https://skm.nib.si), that provides a single, up-to-date entrypoint for plant response investigations.

SKM integrates knowledge on plant molecular interactions and stress specific responses from a wide diversity of sources, combining recent discoveries from journal articles with knowledge already existing in resources such as KEGG (Kanehisa *et al*., 2016), STRING (Szklarczyk *et al*., 2023), MetaCyc (Caspi *et al*., 2016), and AraCyc (Mueller *et al*., 2003). SKM extends other aggregated resources (listed in Supplementary Table 1), including the heterogeneous knowledge graphs of KnetMiner (Hassani-Pak *et al*., 2021), Biomine Explorer (Podpečan *et al*., 2019), and ConsensusPathDB (Herwig *et al*., 2016), in that it allows conversion of biochemical knowledge to diverse mathematical modelling formalisms and integration with multi-omics experiments, besides allowing interactive exploration of current knowledge that is constantly reproducibly updated. SKM is a versatile resource that assists diverse users, from plant researchers to crop breeders, in investigating current knowledge and contextualising new datasets in existing plant research. A number of tools were developed within the SKM environment to support this, and enable efficient linking to complementary tools.

## Results

SKM is a resource combining two knowledge graphs resulting from the integration of dispersed published information on current biochemical knowledge: the **P**lant **S**tress **S**ignalling model (PSS) and the **C**omprehensive **K**nowledge **N**etwork (CKN) of plant molecular interactions. SKM enables interactive exploration of its contents, and represents a basis for diverse systems biology modelling approaches, from network analysis to dynamical modelling.

### The Plant Stress Signalling model (PSS)

PSS is an ongoing endeavour to assemble an accurate and detailed mechanistic model of plant stress signalling by extracting validated molecular interactions from published resources (Miljkovic *et al*., 2012; Ramšak *et al*., 2018). Currently PSS covers the complete stress response cascade within the plant cell (Fig. 1), initiating with abiotic (heat, drought, and waterlogging) and biotic stressors (extracellular pathogens, intracellular pathogens, and necrotrophs; Layer 1). Perception of these stressors through diverse receptors (Layer 2) initiates Ca2+, ROS, and MAPK signalling cascades, as well as phytohormone biosynthesis and signalling pathways (Layer 3). These translate perception into a cellular response, resulting in activation of processes which execute protection against stress (Layer 4). Within and across these layers, relevant transcriptional (transcription factors known to act downstream of phytohormones) and posttranscriptional (e.g. smallRNA-transcript regulation known to participate in stress signalling) regulation is included. To capture the relations between stress responses and growth and development, PSS also contains the major known regulators of growth (Target Of Rapamycin (TOR) signalling) all hormonal signalling pathways and major primary metabolism processes. Finally, tuberisation signalling from potato is included as an example for evaluating potential impact on crop yields.

**Figure 1.**
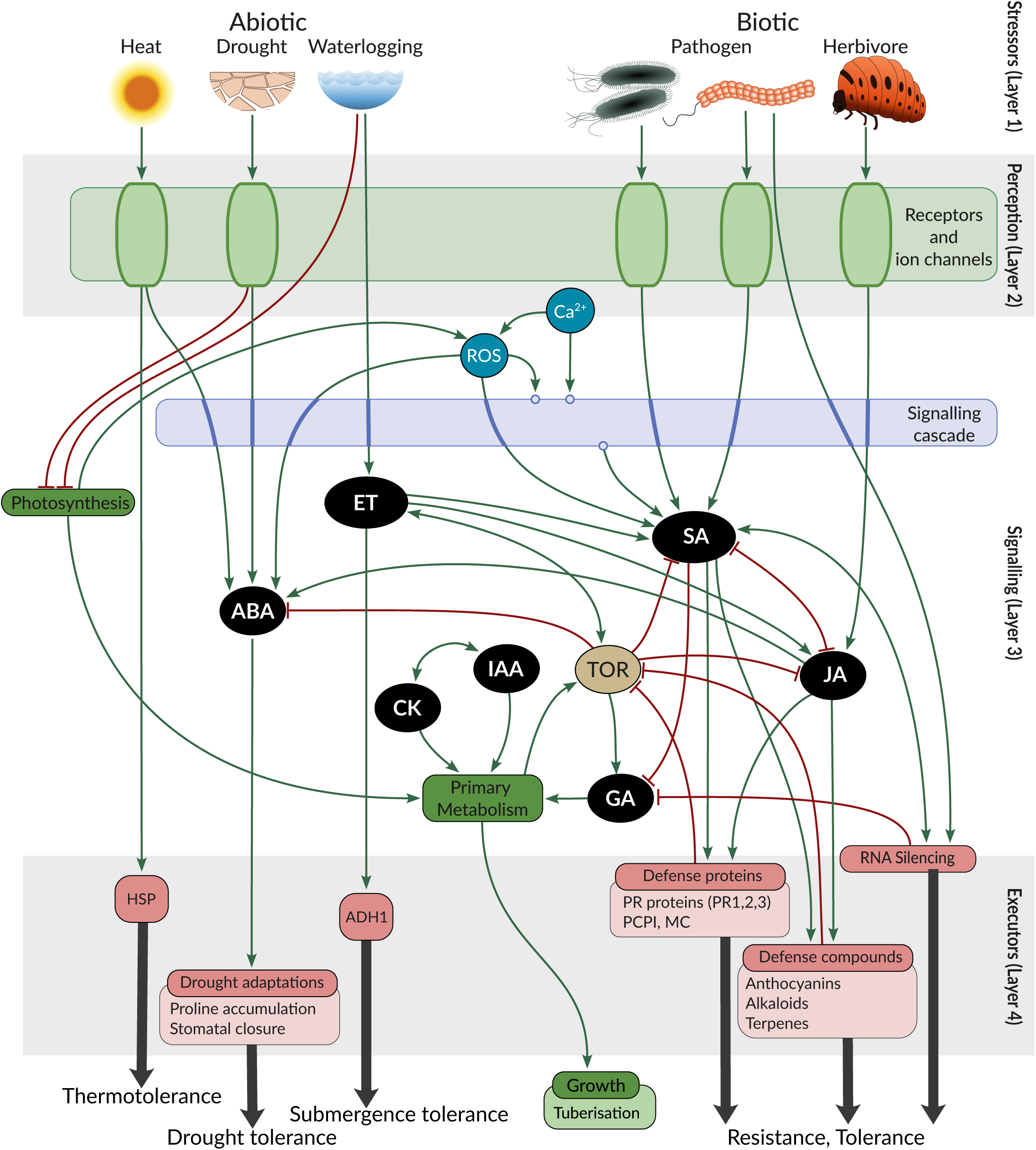
Contents of the Plant Stress Signalling model (PSS) represented as conceptual layers. From top to bottom: stressors (**Layer 1**) acting on the plant are first perceived (**Layer 2**), resulting in a signalling (**Layer 3**) cascade, that leads to plant defence and/or adaptive changes in the form of executor molecules and processes (**Layer 4**, examples listed below each group). ABA: Abscisic Acid; *ADH1*: Alcohol Dehydrogenase 1; CK: Cytokinin; ET: Ethylene; GA: Gibberellic Acid; *HSP*: Heat Shock Protein; IAA: Indole-3-acetic acid (Auxin); JA: Jasmonic Acid; *MC*: Multicystatin, *PCPI*: Potato Cysteine Proteinase Inhibitor; *PR*: Pathogenesis Related; ROS: Reactive Oxygen Species; SA: Salicylic Acid; *TOR*: Target Of Rapamycin.

PSS is primarily based on the model plant Arabidopsis (*Arabidopsis thaliana*), and also contains pertinent information from several crop species, most comprehensively potato (*Solanum tuberosum*). PSS currently includes 1,425 entities and 543 reactions, a substantial update from the preceding model of 212 entities and 112 reactions (Ramšak *et al*., 2018). PSS entities include genes and gene products (proteins, transcripts, smallRNAs), complexes, metabolites, and triggers of plant stress. Genetic redundancy (Cusack *et al*., 2021) is incorporated using the concept of functional clusters – groups of genes (possibly across species) that are known to mediate the same function(s). Interactions between these entities include protein-DNA (e.g. transcriptional regulation), smallRNA-transcript, protein-protein interactions, as well as enzymatic catalysis and transport reactions. The majority of these interactions were compiled from peer-reviewed manuscripts with targeted experimental methodology, giving them a high degree of confidence. PSS also contains relevant signalling associated pathways from KEGG (Kanehisa *et al*., 2016) and AraCyc (Mueller *et al*., 2003).

### The Comprehensive Knowledge Network (CKN)

Complementary to PSS, CKN is a large-scale condition-agnostic assembly of current knowledge, offering broader insights into not only stress signalling, but also any other plant process. CKN is a network of experimentally observed physical interactions between molecular entities, encompassing protein-DNA interactions, interactions of smallRNA with transcripts, post-translational modifications, and protein-protein interactions (Table 1) in Arabidopsis. Here, we present an update to the previous version with 20,012 entities and 70,091 interactions (Ramšak *et al*., 2018), to the current version which provides 30% more entities (26,234 entities) and an almost 7-fold increase in the number of molecular interactions (488,390 unique interactions, Table 1). The entities in CKN include 24,829 genes, out of 38,202 registered in Araport11 (Cheng *et al*., 2017).

**Table 1:** Counts of unique CKN interactions by type and reliability ranking. Rank meanings: 0 – manually curated interactions from PSS, 1 – literature curated interactions detected using multiple complementary (mostly targeted) experimental methods (e.g. luciferase reporter assay, co-immunoprecipitation, enzymatic assays), 2 – interactions detected solely using high-throughput technologies (e.g. high-throughput yeast two-hybrid, chromatin immunoprecipitation sequencing, degradome sequencing), 3 – interactions extracted from literature (co-citation, excluding text mining) or predicted *in silico* and additionally validated with data, 4 – interactions predicted using purely *in silico* binding prediction algorithms. See Supplementary Table 2 for a detailed list of sources.

|  |  | Number of resources | Rank |  |  |  |  | Total |
| --- | --- | --- | --- | --- | --- | --- | --- | --- |
|  |  |  | 0 | 1 | 2 | 3 | 4 |  |
| Interaction type | binding | 13 | 650 | 24,054 | 30,442 | 343,401 | 31,253 | 429,800 |
|  | transcription factor regulation | 9 | 480 | 1,442 | 8,567 | 174 | 11,869 | 22,532 |
|  | small RNA interactions | 3 | - | 48 | 41 | 34,059 | - | 34,148 |
|  | post-translational modification | 2 | 754 | 393 | 192 | - | - | 1,339 |
|  | other<sup>a</sup> | 1 | 571 | - | - | - | - | 571 |
| Total |  | 25<sup>b</sup> | 2,455<sup>c</sup> | 25,937 | 39,243 | 377,634 | 43,122 | 488,390 |

<sup>a</sup> Includes interactions from PSS that do not fall into the previous categories.
<sup>b</sup> Some resources contain multiple interaction types.
<sup>c</sup> Includes interactions expanded from 335 PSS functional clusters to 2253 individual genes.

During the update, only STRING was found to be altered since 2018 (updated to v11.5 in 2021), and thus re-integrated. Additionally, nine novel sources of information were added, bringing the total number of sources CKN integrates to 25 (Supplementary Table 2). Interactions are annotated with the interaction type and whether the interaction has directionality (e.g. undirected binding vs transcription factor regulation). A ranking system for the interaction reliability (Table 1 legend), allows researchers to evaluate how biologically credible and relevant individual interactions are. CKN includes all relevant reactions from PSS to allow for a direct comparison of results obtained through both networks.

### SKM environment and features

To enable accessibility and exploitation of the resources within SKM we have developed an encompassing environment (Fig. 2). The main features include content exploration and visualisation, access to various export formats, and the ability to contribute improvements based on novel biological knowledge. The SKM webpage is publicly available at https://skm.nib.si/.

**Figure 2.**
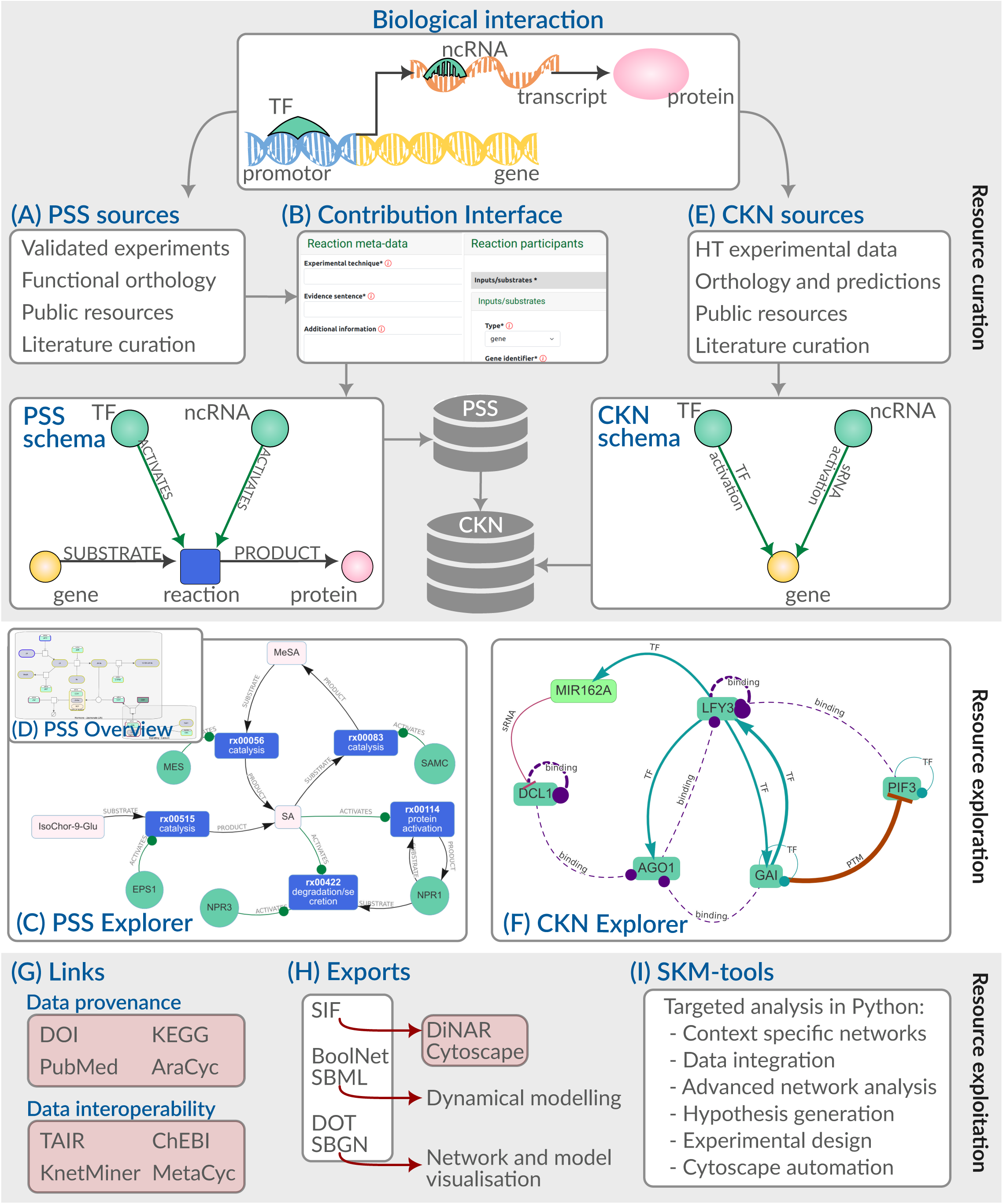
Stress Knowledge Map environment and features. New validated biological interactions (e.g. transcriptional and translational regulation of a target gene) from various **sources (A)** can be added to PSS through the guided contribution interface **(B)**, and are consolidated according to the PSS schema. The contents of PSS can be explored through interactive search and visualisation provided by both the PSS Explorer **(C)** and the PSS overview in Newt **(D)**. Correspondingly, sources for CKN interactions **(E)** are integrated and consolidated to the CKN schema through batch scripts, and are accessible for exploration through the CKN Explorer **(F)** which provides interactive search and visualisation of CKN interactions. Data provenance and interoperability links **(G)** provide context for SKM contents. Exports of PSS and CKN **(H)** enable various additional analysis and modelling approaches, including through the Python functions provided in the SKM-tools resource **(I)**. Links to specific external resources and tools are highlighted in red. HT – high-throughput; PSS – Plant Stress Signalling network; CKN – Comprehensive Knowledge Network; TF – transcription factor; ncRNA – non-coding ribonucleic acid; DOT/SBGN/SBML/SIF – Systems Biology data formats, see Table 3 for details.

**Table 3:** Supported exports of SKM knowledge graphs.

| Format | Description | Available for |
| --- | --- | --- |
| SBGN-ML | Systems Biology Graphical Notation XML format, enabling graphical visualisation of models (Bergmann *et al.*, 2020). | PSS |
| SBML | Systems Biology Markup Language XML format, enabling mechanistic modelling (Keating *et al.*, 2020). | PSS |
| DOT | Graph description language, compatible with Graphviz applications (Gansner and North, 2000) (graphviz.org). | PSS |
| SIF/LGL | Simple Interaction Format/Large Graph Format, compatible with Cytoscape (Shannon *et al.*, 2003) and DiNAR (Zagorščak *et al.*, 2018). | PSS, CKN |
| boolnet | Boolean network format for logical modelling compatible with pyboolnet (Klarner *et al.*, 2017), and BoolNet (Müssel *et al.*, 2010) among others. | PSS |

#### Exploration

SKM implements a number of options for the exploration of its contents, including interactive network visualisations of both PSS (PSS Explorer, Fig. 2C) and CKN (CKN Explorer, Fig. 2F), offering neighbourhood extraction of selected entities, shortest path detection between multiple entities of interest, and on the fly exports. Both Explorers provide direct references to the object provenance, as well as links for the corresponding Arabidopsis genes within KnetMiner knowledge base (Hassani-Pak *et al*., 2021), providing even broader context. An additional visualisation of the complete PSS model, showing biological pathways, is available in the Newt Viewer (Fig. 2D). A separate search interface utilising internal and external database identifiers (e.g. DOI, KEGG) is also available for PSS.

#### Modelling and analysis support

PSS is available for download in a number of domain standard formats (Fig. 2H; summarised in Table 3) enabling further visualisations, analysis, and dynamical modelling. A suite of tools implemented in Python (SKM-tools, Fig. 2I) was developed to support additional network analysis of CKN and PSS (described in Table 4).

**Table 4:** Features of SKM-tools.

| Functionality | Description |
| --- | --- |
| Load | Directly create networkX (Hagberg *et al.*, 2008) graph objects for PSS or CKN, thus providing access to the multitude of graph analysis and graph operations available in the library. |
| Tissue specificity node filter | For PSS and CKN, filter on node type or node origin (plant or foreign), and additionally for CKN filter nodes based on tissue specificity, creating a network specific to the biological question at hand. |
| Edge reliability filter | Filter CKN edges by rank, removing less reliable edges as the situation requires. |
| Network analysis | Standard node based analysis approaches, such as neighbourhood extraction (identifying the immediate interactors of a node) and shortest path analysis (identifying directed or undirected paths between source and target nodes of interest). |
| CUT-tool | CUT-tool provides information on which genes are needed to be perturbed (knock-out, knock-down or overexpress) in order to modulate the response of the network. |
| Cytoscape Automation | Loading of networks and subnetworks into Cytoscape(Otasek *et al.*, 2019). Functionalities include providing default styling, node, edge, and path highlighting, network layout from coordinates, and pdf exporters. |
| Multi-omics data visualisation | Import of multi-omics experimental data tables (e.g. logFC and p-values) as context to the networks, and functionality to visualise experimental data associated with nodes in the network, through rendering of PNG's (e.g. heatmaps) in the Cytoscape view. |
| Link to DiNAR | Instructions for the use of CKN or PSS as the prior knowledge network for integration and visualisation of multiple condition high-throughput data in the DiNAR application (Zagorščak *et al.*, 2018). |

#### Extending and improving SKM

The contribution interface of PSS allows for constant updates based on novel discoveries (Fig. 2B). Registered users can add new entities and interactions to PSS through guided steps, and expert curators are able to make corrections. For major updates to PSS, a batch upload option is also available. The contribution interface automatically retrieves GoMapMan (Ramšak *et al*., 2014) gene descriptions and short names, as well as article metadata via DOI or PubMed ID, simplifying the contribution process.

#### FAIRness

SKM has been developed with the FAIR principles (Findable, Accessible, Interoperable, and Reusable) (Wilkinson *et al*., 2016) at the forefront. SKM is indexed in FAIDARE (FAIR Data-finder for Agronomic Research; https://urgi.versailles.inra.fr/faidare/search?db=SKM), listed in both bio.tools (https://bio.tools/skm) and FAIRsharing.org (https://fairsharing.org/4524), and registered at identifiers.org (https://registry.identifiers.org/registry/skm). Aside from the downloads, a GraphQL endpoint is available for programmatic access to PSS. SKM also utilises stable reaction and functional cluster identifiers. Data provenance is maintained by storing links to input data through DOIs and external database references (Fig 2G).

### Case studies

To showcase the benefits of SKM, we present two case studies utilising SKM for contextualisation of experimental results within prior knowledge networks. The first case study concerns jasmonates (JA) and salicylic acid (SA) interference with abscisic acid (ABA)-mediated activation of RESPONSIVE TO DESICCATION 29 (*RD29*) transcription, and the second a proteomics analysis of Ca^2+^-dependent redox responses.

#### Case study 1: Interaction of ABA, JA, and SA in the activation of *RD29* transcription

In Arabidopsis, the *RESPONSIVE TO DESICCATION 29 A* gene (At*RD29A*) plays a pivotal role in stress acclimation (Baker *et al*., 1994) and is transcriptionally regulated via several promoter elements, including the ABA responsive binding motif ABRE (ACGTG), located close to the transcription initiation site. The 1 kbp upstream region of the potato St*RD29* transcription initiation site also contains the ABA-responsive binding motif ABRE, and several other abiotic stress responsive binding elements (Supplementary Figure 1).

Treatment of leaf discs from tobacco plants transiently transformed with pSt*RD29::fluc* and transgenic potato plants (cv. Désirée) carrying the pSt*RD29::mScarlet-I* (Supplementary Figure 2) construct showed that pSt*RD29* activity was strongly induced by ABA, and reached its highest amplitude after approximately four hours in the ABA solution (Fig. 3A). Treatments with either jasmonate (JA) or salicylic acid (SA) alone did not lead to an increase in pSt*RD29* activity. However, combined treatments of ABA with JA or ABA with SA attenuated the ABA induced activation of pSt*RD29*, indicating a negative impact of both these phytohormones on ABA dependent St*RD29* transcription (Fig. 3A). We subsequently constructed transgenic potato plants (cv. Désirée) carrying the pSt*RD29::fluc* construct to confirm the negative impact of MeJA and SA on the ABA responsive promoter activity *in planta* (Fig. 3B). The impact of MeJA on the ABA activation of both *RD29* was further analysed in potato and Arabidopsis by RT-qPCR. The data revealed that both species display an attenuation of the ABA induction of *RD29A/RD29* by jasmonates (Fig. 3C).

**Figure 3:**
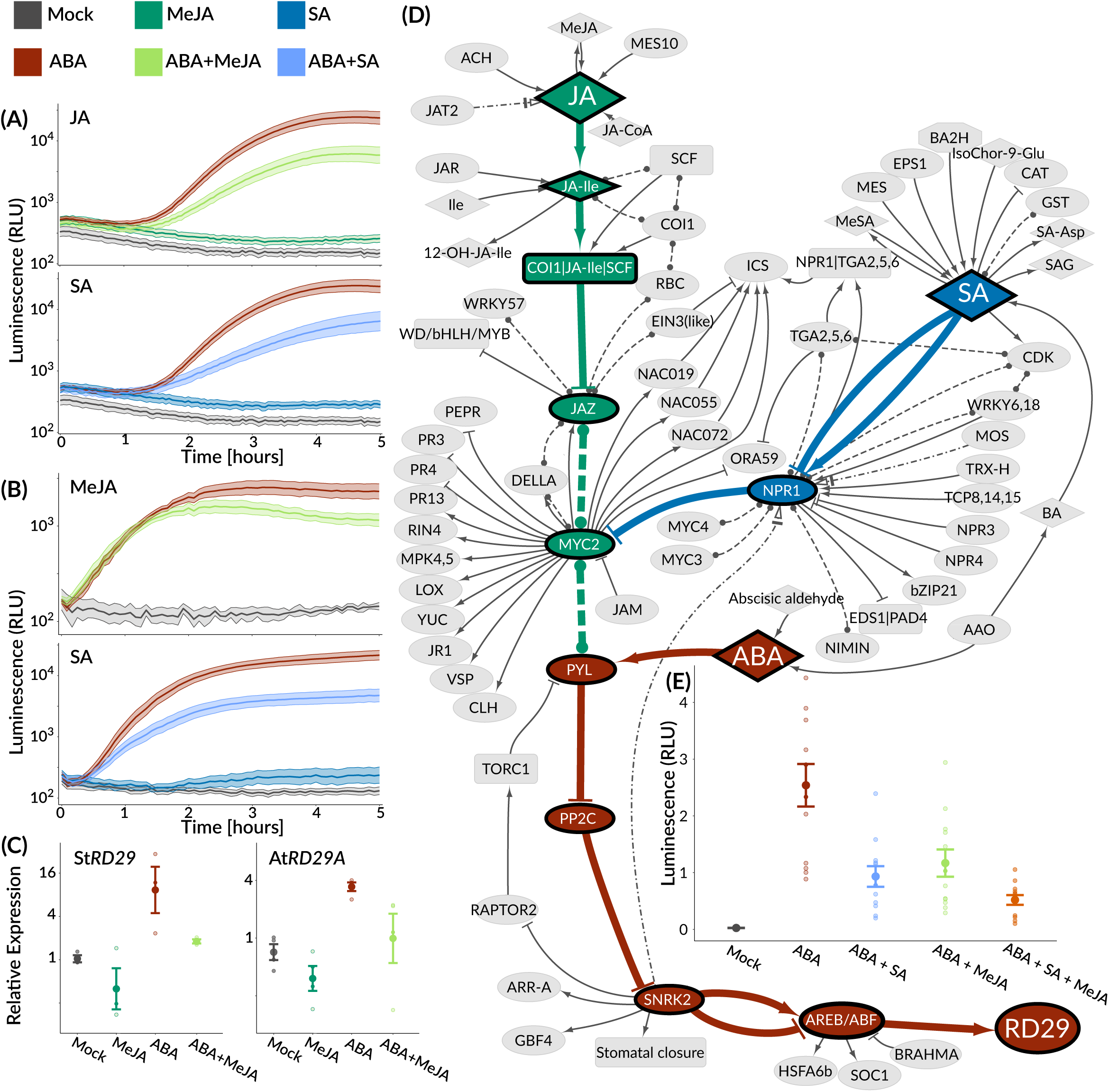
Elucidating connections from JA and SA to ABA-mediated regulation of RD29 expression in potato. **(A/B)** Expression of firefly luciferase driven by the St*RD29* promoter (pSt*RD29*::fluc) in transiently transformed tobacco leaves (A) and transgenic potato leaves (B). Luciferase activity was analysed in response to single and combined phytohormone treatments as indicated (50 µM MeJA, 50 µM ABA and 50 µM SA). Values are shown as mean ± SE. (A) Relative transcript abundance of St*RD29* (left panel) and At*RD29A* (right panel) six hours after application of 50 µM ABA, 50 µM MeJA or combination of both, analysed by RT-qPCR. Bars represent mean values ± SE of 3-4 independent biological replicates. (B) PSS node-induced subnetwork of shortest paths and immediate neighbours. Paths are directed from the hormones (source) to *RD29* (target). Nodes and edges are coloured by the path source: ABA (brown), JA (green), and SA (blue). Edges to first neighbours, edges not on the directed shortest paths, and shared neighbourhood nodes are indicated in grey. Solid edges indicate activation (arrow head) or inhibition (T head), dashed edges represent binding, and dot-dash edges transport. The explorable networks for case study 1 are provided in Supplementary Data 1. (C) Validation of the hypothesis presented in (D). Concentrations of hormones are 50 µM ABA, 15 µM MeJA, and 30 µM SA. Luciferase activity at 5 hours shown (see Supplementary Table 3 for complete response curve). The results show SA and jasmonates indeed act synergistically on attenuation of ABA signalling, as the addition of SA and jasmonates has a stronger effect than the addition of each hormone individually.

We first tried to explain the observed impact of jasmonates and SA on ABA-dependent *RD29* activation through motif analysis of the promoter, but no SA or JA signalling related motifs were identified in the potato promoter sequence (Supplementary Figure 1). Thus we hypothesised that the signalling pathways interact upstream from actual transcriptional activation. Due to the complexity of several phytohormone pathway interactions, this is a good case study for the hormone-centric and expert curated PSS model. We performed a triple shortest path analysis analysis to identify potential mechanisms of studied crosstalk. The analysis revealed an intersection of JA signalling with the ABA pathway through a protein-protein interaction of the JA-responsive MYC-like transcription factor 2 (MYC2) with the ABA receptor PYRABACTIN RESISTANCE LIKE 6 (PYL6; Fig. 3D). This reaction entry (rx00459) is based on experimental *in vitro* and *in vivo* interaction studies of PYL6 and MYC2 in Arabidopsis (Aleman *et al*., 2016). It could be conceived that this interaction depletes PYL, thereby limiting ABA perception(Aleman *et al*., 2016), which could explain lower activation of the ABA pathway in the presence of jasmonates. The SA pathway was found to converge with the ABA pathway through the JA pathway with a protein-protein interaction between the SA receptor NPR1 and MYC2 (rx00432) (Nomoto *et al*., 2021) and this might influence the interaction of MYC with PYL. To verify the hypothesis of direct synergism between JA and SA in the attenuation of the ABA response, we performed titration experiments of combined JA and SA treatment on ABA-dependent St*RD29* induction which was confirmed (Fig. 3E, Supplementary Table 3).

#### Case study 2: The impact of Ca^2+^ channel inhibitor LaCl_3_ on proteome-wide peroxide responses

Secondary messengers, such as Ca^2+^ and H_2_O_2_, are important in the translation of many perceived environmental changes towards a cellular response (Kudla *et al*., 2010; Pirayesh *et al*., 2021). It is still a challenge to disentangle and understand the principles of specificity and information flow in such networks. Lanthanide ions are known to block anion channels and inhibit the flux of Ca^2+^ across the plasma membrane (Knight *et al*., 1992; Tracy *et al*., 2008). Thus, they can be used to identify Ca^2+^-dependent plant responses. H_2_O_2_ is known to induce Ca^2+^ transients (Rentel and Knight, 2004). In this case study, we analysed the proteome of Arabidopsis rosettes treated with either H_2_O_2_ or a combination of H_2_O_2_ and LaCl_3_ to identify the components of H_2_O_2_ signalling that are Ca^2+^-dependent. We initially identified 119 proteins that showed significantly changed abundances in response to H_2_O_2_ compared to mock treatment after 10 or 30 min of treatment. Out of these, 49 proteins did not significantly respond in the same manner upon pretreatment with LaCl_3_ (Supplementary Table 4), indicating that a significant number of H_2_O_2_ induced changes in protein abundance required a Ca^2+^ signal (Ca^2+^-dependent redox-responsive proteins).

In the quest to identify mechanistic explanations behind these results, CKN provides a universal resource for large-scale hypothesis generation. The largest connected component of CKN contains 98% of the nodes and 99% of the edges, indicating its high connectivity, thus the analysis was performed on this part of CKN only. Using CKN pre-filtered to only leaf-expressed genes, we searched for directed shortest paths from known Ca^2+^ signalling related proteins (source set) to the Ca^2+^-dependent redox-responsive proteins identified by the proteomics approach (target set). The final source set of 53 genes included mainly calmodulins, Ca^2+^-dependent protein kinases, and calcineurin B-like proteins CBLs (Supplementary Table 4). Of the 49 Ca^2+^-dependent redox-responsive target proteins, 41 were present in CKN. All of these proteins could either be connected to the source set of Ca^2+^ signalling related proteins directly or through an up to 4-step pathway (Fig. 4A), or were in the source set themselves. Combining all the detected shortest paths (all sources to all targets) into a single network (Fig. 4A) revealed major network hubs – connected to multiple known Ca^2+^ signalling genes and potentially regulating multiple targets.

**Figure 4:**
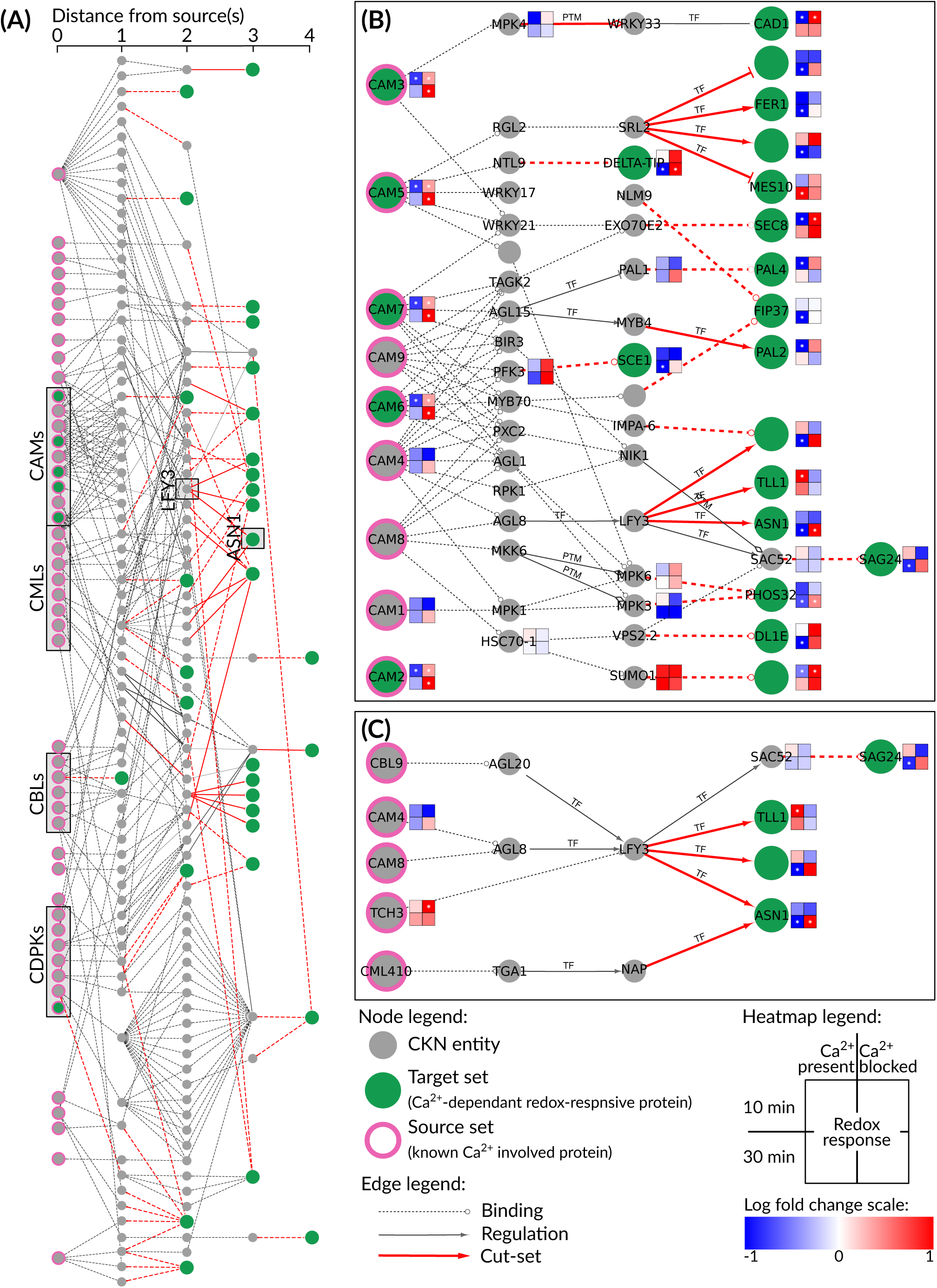
Deciphering the Ca^2+^ dependent network in peroxide signalling. **(A)** All shortest paths identified in CKN leading from known Ca^2+^ related proteins (sources - pink bordered nodes) to Ca^2+^- dependent redox-responsive proteins identified by proteomics (targets - green filled nodes) using rank 0, rank 1, and rank 2 edges (as described in Table 1 legend), merged into a single network. The excerpts show **(B)** a subnetwork with a focus on calmodulins, and **(C)** a subnetwork with a focus on LFY3 and ASN1. Solid edges with arrowheads indicate directed, regulatory interactions (see Table 1), while dashed edges indicate undirected binding. Red edges are part of the merged cut-set. Nodes with proteomics measurements are annotated with a heatmap indicating change in protein abundance after 10 min (top row), after 30 min (bottom row) between H_2_O_2_ and mock treated samples (left column) and between Ca^2+^ blocker treatment and H_2_O_2_ and Ca^2+^ blocker treatment (right column). Significant changes in abundance are marked with an asterisk in the centre of the square. Red – increase in treatment compared to control, blue – decrease in treatment compared to control. Nodes are labelled with their short name, if it exists. The complete explorable networks are provided in Supplementary Data File 1, and all source and target nodes are listed in Supplementary Table 4.

The analysis, for example, revealed an intricate network of calmodulins-dependent regulation of downstream targets in Arabidopsis (CAM2,3,5,6,7, Fig. 4B), Another example of such a hub is Floricaula/leafy-like transcription factor 3 (*LFY3*) shown in Fig. 4C, which integrates paths originating from four source nodes, and in turn potentially regulates four downstream targets.

The next step in the analysis would be confirmation of the identified mechanisms with functional analysis experiments, e.g. to perform knock-out experiment(s) and confirm the role of the proposed regulatory network. The design of such experiments is however not always trivial, thus we designed the CUT-tool within SKM-tools, to aid experimentalists. This analysis reveals the minimum interactions that are necessary to be severed (“cut-set”) in order to separate the upstream regulators from the downstream targets. The cut-set to disrupt the regulation of all targets are shown in Fig 4A. As an example, the cut-set of one target, glutamine-dependent asparagine synthase 1 (ASN1) is shown in Fig. 4C, revealing that de-regulation of ASN1 would require the knockout of both LFY3 and ARABIDOPSIS NAC DOMAIN CONTAINING PROTEIN 29 (NAP).

## Discussion

Plant stress signalling pathways are connected by synergistic and antagonistic interactions in a complex network that checks and balances the plant’s response to their environment and its growth/development (Eckardt, 2015; Bittner *et al*., 2022). To understand the functioning of these complex processes, novel approaches are required. Knowledge graphs, such as those provided by SKM, provide powerful and accessible tools to integrate and simplify interpretations within curated published knowledge, as well as providing a basis of a plant digital twin, and all the advantages of *in silico* simulation experiments it enables. A number of tools were developed within the SKM environment to support this, and also enable efficient linking to complementary tools.

To showcase the applicability of SKM, we investigated two distinct experimental datasets. In the first, our experiments showed evidence that jasmonate and SA treatment attenuates ABA activated transcription of *RD29* in both the crop plant potato and the model plant Arabidopsis through hormonal signalling cross-talk (Fig. 3). A manual attempt to extract known information on crosstalk between ABA and JA with a search in PubMed (*(JA OR jasmon*) AND (ABA OR abscisic) AND (plant)*) resulted in over 2,000 published items. With the wealth of data generated these days, it would be laborious for an individual researcher to perform a thorough literature survey, while interrogation of SKM provided a mechanistic hypothesis that explains the experimental results within hours. The hypothesis was experimentally confirmed and gives the explanation for the synergistic action of jasmonates and SA that is sometimes argued for in literature (Mur *et al*., 2006; Zhang *et al*., 2020). Although knowledge compiled in SKM is predominately based on Arabidopsis, this use case clearly shows its applicability in other species. Through orthology tools such as PLAZA (Van Bel *et al*., 2022), the knowledge graphs in SKM can be translated to other species, as was done for the previous version of CKN to *Prunus persica* (Foix *et al*., 2021), *Solanum tuberosum* (Ramšak *et al*., 2018), and *Nicotiana benthamiana* (Juteršek *et al*., 2022). This way, canonical principles of plant signalling networks can be assessed across species.

Our second case study showed that SKM is not only helpful in revealing mechanisms in complex pathways for a single target, but also can be used to identify regulators using a large number of targets, as is commonly the case with interpretation of large omics datasets. Using network analyses, arguably the simplest qualitative modelling approach, we identified hubs involved in complex redox - Ca^2+^ signalling interconnectedness. By identifying connections from known Ca^2+^ related proteins to our experimentally derived target list, we were able to prioritise certain processes and hypotheses in an informed manner. One of the SKM-tools features, the CUT-tool, was designed to help in the next step of research: validation of generated hypotheses. It allows for the design of complex functional validation experiments (e.g. gene knock-out or overexpression) identifying the genes whose activity should be modulated to achieve a desired effect, taking network redundancy into account. Overall, in both case studies, SKM proved to be a useful generator of potential mechanistic explanations of the observed data.

In agriculture, plant digital twins, as virtual replicas of physical systems, are expected to provide a revolutionary platform for modelling the effect of crop management systems and environmental changes (Pylianidis *et al*., 2021). Digital twins can be used to perform *in silico* experiments that guide or replace lab and field experiments. The detail that digital twins provide, combined with fast computational methodologies, allows for efficient planning of experiments and will thus speed up our understanding of plant functioning and provide information for more effective breeding. Aside from being a tool for the interpretation of experimental data, SKM also provides a starting point for the integration of stress signalling and growth tradeoffs in digital twins.

SKM will be continuously updated, keeping abreast of the latest developments in the field. We believe the integrated knowledge in SKM will help in understanding of plant interactions with the environment, by enabling exploration of knowledge and by supporting diverse mechanistic modelling approaches. This is of interest to the wider plant scientific community, enabling the informed design of experiments and, in the long term, contributing to the breeding of improved varieties and precision agriculture.

## Methods

### PSS construction

From the predecessor model (“PIS-v2”, Ramšak *et al*., 2018), numerous improvements, additions, and reformulations were carried out, resulting in the current PSS. In addition to intracellular pathogens (potyviruses), we extended PSS to also contain perception of extracellular pathogens (*Pseudomonas* sp.) and insect pests, as well as heat, drought, and waterlogging stress. Downstream of perception, PSS now includes Ca^2+^ signalling, ROS signalling, the MAPK signalling cascade, as well as the synthesis and signalling of all major phytohormones. We also added the synthesis of actuator molecules and processes, as well as known regulators of growth and major processes leading to growth.

PSS is implemented as a Neo4j graph database. The types of nodes and edges (relationships) in the database are summarised in Supplementary Table 5. Genes and gene products are represented by functional cluster nodes, including protein and noncoding RNA nodes. Functional clusters allow for the representation of genetic redundancy. These groups were defined using sequence similarity between genes (orthologues and paralogues) and experimental data that confirmed functional overlap. The functional cluster concept includes groupings of enzyme coding genes (similarly to the E.C. number system), as well as genes involved in transcriptional and translational regulation. Groups of metabolites with the same biological function are also represented as metabolite families. Nodes also include more abstract entities, such as known but unidentified gene products and plant processes. Finally, foreign entities, such as biotic or abiotic stressors are also included as nodes.

In addition to biological entities, molecular interactions are also represented by nodes in PSS, and are categorised into ten formal reaction types (e.g. protein activation or catalysis, Supplementary Table 5). Reaction participant nodes are connected to the reaction nodes by relationships, with the type of relationship representing the role of the participant (e.g. SUBSTRATE, ACTIVATES), as demonstrated in Fig. 2B. These relationships are annotated with the subcellular location and the form of the participant when involved in the reaction (e.g. ‘cytoplasm’ or ‘nucleus’ and ‘gene’ or ‘protein’).

Where applicable, nodes are annotated with their provenance (e.g. a DOI) and additional information such as biological pathways, gene identifiers, descriptions and annotations (TAIR (Berardini *et al*., 2015), GoMapMan (Ramšak *et al*., 2014)), references to external resources (DOI, PubMed, KEGG (Kanehisa *et al*., 2016), MetaCyc (Caspi *et al*., 2016), AraCyc (Mueller *et al*., 2003), and ChEBI (Hastings *et al*., 2016)), and explanatory statements (such as a quote from the article and the experimental techniques used in the original experiments).

All updates to PSS are immediately available in the various interfaces and all download formats. A frozen version (PSS v1.0.0) is also available in all export formats and additionally, a database dump with detailed deployment instructions can be accessed at GitHub (https://github.com/NIB-SI/skm-neo4j). All sources and resources used to create PSS v1.0.0 are available in Supplementary Table 6.

PSS is available in a number of systems biology standard formats, including SBML (using libSBML (Bornstein *et al*., 2008)), SBGN (using libSBGN (König, 2020) and pySBGN (Podpečan, 2023) libraries), DOT (using pygraphviz (Aric Hagberg *et al*.) and pydot (Sebastian Kalinowski *et al*., 2023)), and a Boolean formulation in boolnet format. SKM also supplies several generalised formats of PSS in SIF/TSV format, allowing multiple formulations of the network model.

### CKN construction

The second edition of the comprehensive knowledge network (CKN-v2) was created by merging pairwise interactions from 25 public resources (details in Supplementary Table 2). Additional filtering was performed on the STRING v11.5 network (Szklarczyk *et al*., 2023), where the requirement was to only include physical interactions, confirmed by experimental data or existence in a database. As Table 2 summarises, five reliability ranks were designed to describe the reliability of the interactions, across the diversity of the various sources. All interactions were then integrated, resulting in a single network of 574,538 interactions. The network was subsequently condensed by collapsing multiple interactions of the same type between a pair of interactors into a single edge. In this process, the highest ranked interaction took precedence to define the interaction type, but all sources that contain any interaction between the pair were retained in the edge attributes.

All gene loci nodes were annotated using Araport11 (Cheng *et al*., 2017) downloaded from TAIR in June 2023 (Berardini *et al*., 2015). Gene loci that have been merged or made obsolete were renamed or removed respectively. Genes are also annotated with Plant Ontology annotations from TAIR (Berardini *et al*., 2015) (based on gene expression patterns reported in publications), enabling the extraction of tissue specific interaction networks.

CKN-v2 is available as part of the SKM application and on the downloads page (https://skm.nib.si/downloads/).

### SKM Environment

The SKM web application is implemented in Python using the microframework Flask. The interactive visualisations of PSS and CKN are based on Biomine Explorer (Podpečan *et al*., 2019), implemented using vis.js and open-source Python libraries (including networkX (Hagberg *et al*., 2008) and graph-tools (Peixoto, 2014)), and are freely available on GitHub at https://github.com/NIB-SI/ckn_viz and https://github.com/NIB-SI/pss_viz respectively. The mechanistic interface to PSS is provided through an instance of the Newt Editor(Balci *et al*., 2021), utilising the SBGN standard.

### SKM-tools

SKM-tools (https://github.com/NIB-SI/skm-tools) is a collection of Python scripts and notebooks, incorporating network analysis and visualisation tools, that facilitates interrogation of CKN and PSS with targeted questions beyond the scope of the web application. Included functionalities are described in Table 4. The tools are developed using the networkX (Hagberg *et al*., 2008) and py4cytoscape (Keiichiro Ono *et al*.) libraries.

The CUT-tool utilises the max-flow min-cut (Edmonds-Karp (Edmonds and Karp, 1972)) algorithm, which determines the minimum edges that are necessary to be severed (“cut set”) in order to separate the upstream sources from downstream targets. A max-flow min-cut analysis of multiple sources to an individual target reveals the minimum cut set to disrupt all signalling to the target. In order to calculate the max-flow min-cut across multiple sources, a dummy node connected with arbitrarily high capacity to all original sources is introduced, and the calculation done using the dummy node as the source.

### Case studies

#### Promoter analysis

Predicted cis-regulatory motifs within the 1kbp promoter sequence of At*RD29A* and St*RD29* were identified via the Atcis-database of the Arabidopsis Gene Regulatory Information Server (AGRIS) (Lichtenberg *et al*., 2009). In addition we used PlantPAN 3.0 (Chow *et al*., 2019) to identify St*RD29* specific motifs which were not previously identified in At*RD29A*.

#### Plant material and growth conditions

*Solanum tuberosum* (cv. Désirée) plants were propagated by cuttings from sterile grown plants. After 7 days of sterile growth on ½ MS-Media (pH 5.7, 2 (w/v) % sucrose) to initiate root growth, plantlets were transferred into single pots filled with soil (9 parts soil, 1 part perligran). *Arabidopsis thaliana* (ecotype Col-0) seeds were directly sown on soil and transferred into single pots after 4-6 days. For all experiments, leaves were used from 18-21 days old plants grown in climatized chambers (20 ± 2 °C) under long-day conditions (16 h light/8 h dark) with a light intensity of 120 μmol photons m^-2^ s^-1^ (Philips TLD 18W alternating 830/840 light colour temperature).

For promoter reporter assays of transiently transformed *N. benthamiana* leaves, seeds were germinated on pProfi-substrate (Gramoflor). Five days after germination, seedlings were separated into 15.5 cm diameter x 12 cm height pots of 15.5 cm diameter x 12 cm height filled with substrate (3 parts profi-substrate, 1 part vermiculite, 1.5 kg osmocote start per m³). Plants were grown in a greenhouse under long day conditions (16h light at 28 °C/8 h dark at 22 °C) at an average light intensity of ∼250 µE and 80% relative humidity.

Soltu.DM.03G017570 was identified as the orthologous locus of Arabidopsis *RD29A* in *S. tuberosum* cultivar DM1-3 using the DM v6.1 database (http://spuddb.uga.edu/). To generate the gene reporter lines in the potato cv. Désirée, 1158 bps of the 5’ UTR directly upstream of the start codon region were amplified by PCR and either the firefly *luciferase* (fluc) or the *mscarletI* (*mScar*) gene in a custom variant of the pBIB Hyg vector carrying a hygromycine resistance for selection in plants. The complete sequences of both vectors including annotations can be found in Supplementary Figure 3. Both constructs were introduced into the potato cultivar Désirée as described previously (Rocha-Sosa *et al*., 1989).

#### Plate-reader based luciferase assays

Agrobacteria carrying the pBIN-StRD29::fluc or pBIN-AtRD29A::fluc plasmid were grown in LB liquid medium supplemented with the respective antibiotics. Overnight cultures were diluted to OD600 = 0.1 with fresh LB medium and grown to OD600 = 0.8. Cells were harvested by centrifugation (22°C, 15 min 4000g) and resuspended in 5% sucrose solution in H_2_O to an OD600 = 0.2. The agrobacteria suspension was infiltrated into leaves #6, #7 and #8 of four week old *N. benthamiana* plants. Care was taken that the *N. benthamiana* plants selected for infiltration and measurement were not suffering an obvious pathogen attack before infiltration, during the transformation period, hormone treatment and measurement. After 48 hours, leaf discs (ø 6 mm) of infiltrated plants were transferred into 96 well plates containing 100 µl buffered MS (5 mM MES, pH 5.8) supplemented with 1 % sucrose (w/v) and incubated for 2 hours under greenhouse growth conditions. Immediately before measurement, luciferin, to a final concentration of 30µM and the hormones, to the final concentration indicated in the text, were added into each respective well. For all combinatorial hormone treatments the different hormones were applied at the same time to the indicated final concentrations. Fluc-luminescence was recorded in a multi-mode microplate reader (TECAN spark multimode microplate reader, Serial number: 2301004717) in a window from 550 nm to 700 nm, for 2 seconds every 5 min for each well. During the measurement period the leaf discs were kept in darkness and at a constant temperature of 22 °C.

For luminescence measurements on *S. tuberosum* St*RD29*::fluc plants, leaf discs (ø 6 mm) were placed in a 96-well plates containing 100 μl of 30 μM luciferin dissolved in ½ MS After 2 hours of preincubation, the solution was replaced by 100 µl of 30 µM luciferin containing various effectors (50 µM ABA, 50 µM MeJA or mix of both) and luminescence was measured every 5 min for up to 12 hours using aTriStar2 lb 492 multimodereader (Berthold Technologies GmbH, Germany). During the measurement period the leaf discs were kept in darkness. All luminescence analysis was performed with at least 5 independent experimental replicates. Luminescence data is available in Supplementary Table 3 and Supplementary Table 7.

#### Transcript analysis

For St*RD29* and At*RD29A* transcript analysis, *S. tuberosum* or *A. thaliana* plants were treated with water (mock), 50 µM ABA, 50 µM MeJA or combination of both for 6 hours in 3-4 independent biological replicates. Total RNA was extracted from 100 mg leaf material using the Gene Matrix Universal RNA Purification Kit (Roboklon, Germany) according to the manufacturer’s instructions. RNA integrity was assessed by agarose electrophoresis and RNA quantity and purity by UV/VIS spectrophotometer (Eppendorf, Germany). For quantitative real-time PCR (qRT-PCR) analysis, RNA was transcribed into cDNA using the RevertAid First Strand cDNA Synthesis Kit (Thermo Scientific, Germany). The reaction was stopped by 5 min incubation at 75 °C.

Where applicable, all primers were designed to span exon–intron borders using QUANTPRIME (Arvidsson *et al*., 2008) (gene identifiers and primer sequences in Supplementary Table 8). qRT-PCR was performed with three technical replicates for each sample in 96 well plates using a CFX96 real-time thermal cycler system (Bio-Rad, Germany). Each reaction contained 1x SYBR-green master mix (Thermo Fisher), 2 ng/µl cDNA and 10 µM each of the respective forward + reverse primer. The specificity of each product was assessed based on the melting curves after 40 cycles of amplification. All transcript levels were normalised against the geometric mean of the transcript abundances of the reference genes *YLS8* and *CYP5* for Arabidopsis and *YLS8* and *ACT7* for potato. Target relative copy numbers were calculated using quantGenius (Baebler *et al*., 2017) (http://quantgenius.nib.si/), provided in Supplementary Table 9.

#### PSS network analysis

We identified the pathway between ABA and *RD29* by querying for all directed shortest paths from ABA to *RD29* in the reaction participant bipartite projection of PSS. We then extracted all directed shortest paths from JA and SA to *RD29* that partially overlapped with the ABA to *RD29* path. For added context to these results, we expanded the network induced by the shortest paths to include the first neighbours of all nodes (Fig. 3E).

Analysis was performed in Python using the networkx (Hagberg *et al*., 2008) library and visualised in Cytoscape (Cline *et al*., 2007) using the py4cytoscape (Keiichiro Ono *et al*.) library. All code is available in the SKM-tools repository (https://github.com/NIB-SI/skm-tools).

#### Proteomic analysis

Complete rosettes of three-week-old *A. thaliana* plants were incubated in 1 mM LaCl_3_ solution or ddH_2_O for 1 hour. Afterwards, plants were transferred into either 20 mM H_2_O_2_ or into ddH_2_O and harvested after 10- and 30-min incubation, respectively. Complete rosettes of 12 plants per treatment were pooled and immediately frozen in liquid nitrogen. Frozen plant material was homogenised using a pre-cooled mortar and pestle and stored at -80 °C. For peptide isolation, 500 mg frozen plant material was mixed with 2 ml lacus-buffer (20 mM Tris pH 7.7, 80 mM NaCl, 0.75 mM EDTA, 1 mM CaCl_2_, 5 mM MgCl_2_, 1 mM DTT, 1/200 mM NaF) containing 4 tablets of protease inhibitor (Roche cOmplete, EDTA-free, Protease inhibitor cocktail tablets) and 10 tablets of phosphatase inhibitor (Roche PhosSTOP™) per 200ml. Samples were incubated for 10 min on ice and subsequently centrifuged at 15.000 g for 10 min at 4 °C. The supernatant was transferred into a new tube, adjusted to 20% (v/v) trichloroacetic acid and incubated overnight at -20 °C. The precipitated samples were stored until preparation for mass-spec analysis.

Samples were centrifuged at 15.000 g, vacuum-dried and eluted in urea lysis buffer (8 M urea, 150 mM NaCl and 40 mM Tris-HCl pH 8). Protein concentration was determined via BCA-assay (Thermo Fisher). In total, 3 mg of protein per sample were first reduced in 5 mM DTT and subsequently alkylated in 15 mM iodoacetamide for 30 min at room temperature in the dark. The alkylated samples were quenched by adding DTT to final concentration of 5 mM and mixed with 30 mg Sera-Mag carboxylate-modified magnetic beads (1:1 ratio of hydrophilic and hydrophobic beads, Cytiva, USA). The peptides attached to the beads were washed four times with 80% (v/v) ethanol and digested in a 30 mM ammonium bicarbonate buffer (pH 8.2) containing 30 µg trypsin (Promega, Wisconsin, USA). Tryptic digestion was performed overnight at 37 °C under constant shaking. The digestion was stopped by the addition of formic acid (end-concentration of 4%). In total, 100 µg of the digested peptides per sample were transferred into a new reaction tube, vacuum-dried and stored at -20 °C until HPLC-MS/MS analysis.

The purified tryptic peptides were dissolved in 0.1% (v/v) formic acid in high purity water. Approximately 1 µg of peptides were separated by an online reversed-phase HPLC (Thermo Scientific Dionex Ultimate 3000 RSLC nano LC system) connected to a benchtop Quadrupole Orbitrap (Q-Exactive Plus) mass spectrometer (Thermo Fisher Scientific). The separation was carried on an Easy-Spray analytical column (PepMap RSLC C18, 2 μm, 100 Å, 75 μm i.d. × 50 cm, Thermo Fisher Scientific) with an integrated emitter, and the column was heated to 55°C. The LC gradient was set to a 140-min gradient method, with a flow rate of 300 nL/min. The LC gradient was set to 5 - 50% buffer B (v/v) [79.9% ACN, 0.1% formic acid, 20% Ultra high purity (MilliQ)] for 125 min, and then to 80% buffer B over 5 min.

LC eluent was introduced into the mass spectrometer through an Easy-Spray ion source (Thermo Scientific), with the emitter operated at 1.9 kV. The mass spectra were measured in positive ion mode applying a top fifteen data-dependent acquisition (DDA). A full mass spectrum was set to 70,000 resolution at m/z 200 [Automatic Gain Control (AGC) target at 1e6, maximum injection time (IT) of 120 ms and a scan range 400-1600 (m/z)]. The MS scan was followed by a MS/MS scan at 17,500 resolution at m/z 200 (AGC target at 5e4, 1.6 m/z isolation window, and maximum IT of 80 ms). For MS/MS fragmentation, normalised collision energy (NCE) for higher energy collisional dissociation (HCD) was set to 27%. Dynamic exclusion was set at 40 s, and unassigned and +1, +7, +8, and > +8 charged precursors were excluded. The intensity threshold was set to 6.3e3, and isotopes were excluded. The analysis was performed with 5 independent experimental replicates for each sample.

#### Peptide identification and quantification

Identities and peptide features were defined by the peptide search engine Adromeda, which was provided by the MaxQuant-software (Version 2.1.3.0, Max Planck Institute of Biochemistry) using standard settings (Tyanova *et al*., 2016b). In detail, trypsin based digestion of the peptides with up to two missing cleavage sites were selected. Methinonine-oxidation as well as N-terminal acetylation was set as variable modifications for peptide identification. In total, up to three potential modification sites per peptide were accepted. The identified peptide sequences were searched and aligned against the Araport11 (Cheng *et al*., 2017) reference protein database. The FDR cut-off for protein identification and side identification was set to 0.01. The minimum peptide length was 7 AA and the maximum length was 40 AA. For each identified protein group, label-free quantitation intensities were calculated and used for further analysis (Supplementary Table 4).

Potential contaminants and reverse sequenced peptides were removed before statistical analysis. Only proteins that were detected in at least three out of five replicates in at least one treatment group were considered for statistical analysis, which was performed using the Perseus (Version 2.0.7.0) (Tyanova *et al*., 2016a). Missing values were replaced by sampling from a normal distribution using the default settings. Protein groups with an absolute fold change of above 1.5 compared to the control and a FDR value below 0.05 were considered as significantly regulated (Supplementary Table 4).

To filter for Ca^2+^-regulated proteins, significantly up (down) regulated proteins in La^3+^ + H_2_O_2_ compared to La^3+^ only treated samples were subtracted from the list of significantly up (down) regulated proteins in H_2_O_2_ treated samples. An additional filtering step was performed to ensure a compelling difference in abundance between the two contrasts. This required that *abs(L_1_ - L_2_) ≥ 1*, where *L_1_* = log fold change for H_2_O_2_ vs mock and *L_2_* = log fold change for La^3+^ + H_2_O_2_ treatment vs La^3+^ only. For each of the protein groups that passed the filters, we extracted all identifiers in the group. For identifiers which occurred in multiple groups, we removed the identifier from the group where it occurred the least.

#### CKN network analysis

For each Ca^2+^-dependent redox-responsive protein group (target), we identified the closest nodes upstream that have a known Ca^2+^ signalling association (source). This was done by identifying all shortest paths in CKN with the source nodes set as all genes with Ca^2+^ signalling related GoMapMan (Ramšak *et al*., 2014) annotations and the target set as the Ca^2+^ dependent H_2_O_2_ responsive peptides. The GoMapMan annotations considered were ‘30.3 - signalling.calcium’, ‘34.21 - transport.calcium’, and ‘34.22 - transport.cyclic nucleotide or calcium regulated channels’. For each target, we kept the source(s) with the shortest paths to the target (the “closest” upstream potential Ca^2+^ interactors). We used the CUT-tool on the merged network to determine the cut set between all the source nodes and each target. The capacity on the edges was set as the edge rank + 1 (highly ranked edges are more likely to be in the cut set).

All source and target nodes are listed in Supplementary Table 4. Analysis was performed in Python using the networkx (Hagberg *et al*., 2008) library and visualised in Cytoscape (Cline *et al*., 2007) using the py4cytoscape (Keiichiro Ono *et al*.) library. All code is available in the SKM-tools repository (https://github.com/NIB-SI/skm-tools).

### Gene identifiers

All genes mentioned in the article are listed with their gene identifiers in Supplementary Table 10.

## Author Contributions

**Software** and **Visualisation** for SKM was done by CB and VP. **Data Curation** of CKN was done by ŽR and CB. **Data Curation** of PSS was performed by CB, ŽR, MZ, ŠB, MP, MK, AŽ, and KG. **Supervision**, **Project administration**, and **Funding acquisition** for SKM development was performed by KG. For case study one **Investigation** was performed by AB, BW, JG, **Formal analysis** by CB and AB, **Data Curation** by MZ and ŠB, **Visualisation** by CB, AB, and MZ. **Supervision** of case study one was performed by UV, MT, KG, and UV, MT provided **Project administration** and **Funding acquisition**. For case study two, **Methodology** was performed by LAS, **Investigation** was performed by AB, BW, AVD, and LAS, **Formal analysis** by CB and AB, **Data Curation** by AB, BW, AVD, and LAS, **Visualisation** by CB and AB. **Supervision** of case study two was performed by UV and KG, and UV provided **Project administration**. **Writing - Original Draft** was performed by CB, AB, ŽR, and KG. All authors took part in **Writing - Review & Editing**.

## Acknowledgements

The authors would like to acknowledge Solana Research GmbH for producing St*RD29*::fluc Désirée transgenic potatoes, and Nelly Braun for pre-selection of the St*RD29* lines. The authors would also like to thank the many additional contributors and curators of PSS: Anna Coll, Barbara Jaklič, Christian Bachem, Christian Schuy, Juan Antonio López-Ráez, Katja Stare, Maria Pozo, Mojca Juteršek, Špela Tomaž, Tim Godec, Tjaša Lukan, Tjaša Mahkovec Povalej, Valentina Levak, Vaňková Radka, Vid Modic, and Maroof Shaikh. Finally, the authors would like to acknowledge Zoran Nikoloski for discussions regarding the CKN analyses.

## Supplementary information

| Name | File name | Description |
| --- | --- | --- |
| Supplementary Table 1 | S01_SupplementaryTable1_ComparativeResources.xlsx | A non-exhaustive list of complementary, comparative, and integrated resources of SKM. |
| Supplementary Table 2 | S02_SupplementaryTable2_CKNv2-sources.xlsx | List of sources of CKN-v2 interactions. |
| Supplementary Figure 1 | S03_SupplementaryFigure1_MotifsAtRD29A-StRD29.pdf | Visualisation of abiotic stress related cis-regulatory binding motifs for *AtRD29A* and *StRD29*. |
| Supplementary Figure 2 | S04_SupplementaryFigure2_ABA-response-of-StRD29.pdf | Microscopic (CLSM) analyses of ABA response of *St-RD29::mScarletI*, showing that ABA activates *St-RD29::mScarletI* in stomata of potato plants. |
| Supplementary Table 3 | S05_SupplementaryTable3_Luminescence-RD29-synergistic.xlsx | Luminescence data for case study 1 showing *StRD29* expression induction by ABA, and validation of the hypothesis of synergistic activity of combinatorial jasmonates and SA in attenuation of expression. |
| Supplementary Data 1 | S06_SupplementaryData1_Case-studies-Cytoscape.cys | Case study 1 (PSS) and Case study 2 (CKN) network analysis results provided in a Cytoscape session. |
| Supplementary Table 4 | S07_SupplementaryTable4_Case-study-2-Proteomics-and-CKN-analysis.xlsx | Case study 2 proteomics data, processed proteomics data, gene descriptions, and CKN network analysis results. |
| Supplementary Table 5 | S08_SupplementaryTable5_PSS-schema.xlsx | PSS database schema description. |
| Supplementary Table 6 | S09_SupplementaryTable6_PSS-sources.xlsx | List of sources of PSS v1.0.0 interactions. |
| Supplementary Table 7 | S10_SupplementaryTable7_Luminescence-RD29.xlsx | Luminescence data for case study 1 showing *StRD29* expression induced by ABA, attenuated by addition of jasmonates or SA. |
| Supplementary Figure 3 | S11_SupplementaryFigure3_Vector-StRD29-fluc-and-StRD29-mScarletI.pdf | Visualisation of the features of Vector pBibHyg carrying *StRD29::fluc* and *StRD29::mScarletI*. |
| Supplementary Table 8 | S12_SupplementaryTable8_Primer-sequences.xlsx | Gene identifiers and primer sequences for transcript analysis. |
| Supplementary Table 9 | S13_SupplementaryTable9_RD29-qPCR.xlsx | Relative gene expression of *RD29* in potato and *Arabidopsis* after treatment with ABA, JA or their combination. |
| Supplementary Table 10 | S14_SupplementaryTable10_Gene-identifiers.xlsx | Genes and gene identifiers mentioned in the article. |

## References

Aleman, F., Yazaki, J., Lee, M., Takahashi, Y., Kim, A. Y., Li, Z., Kinoshita, T., Ecker, J. R., and Schroeder, J. I. (2016). An ABA-increased interaction of the PYL6 ABA receptor with MYC2 Transcription Factor: A putative link of ABA and JA signaling. Sci. Rep. 6:28941.

Aric Hagberg, Dan Schult, and Manos Renieris PyGraphviz. https://pygraphviz.github.io/

Arvidsson, S., Kwasniewski, M., Riaño-Pachón, D. M., and Mueller-Roeber, B. (2008). QuantPrime – a flexible tool for reliable high-throughput primer design for quantitative PCR. BMC Bioinformatics 9:465.

Baebler, Š., Svalina, M., Petek, M., Stare, K., Rotter, A., Pompe-Novak, M., and Gruden, K. (2017). quantGenius: implementation of a decision support system for qPCR-based gene quantification. BMC Bioinformatics 18:276.

Baker, S. S., Wilhelm, K. S., and Thomashow, M. F. (1994). The 5′-region of Arabidopsis thaliana cor15a has cis-acting elements that confer cold-, drought- and ABA-regulated gene expression. Plant Mol. Biol. 24:701–713.

Balci, H., Siper, M. C., Saleh, N., Safarli, I., Roy, L., Kilicarslan, M., Ozaydin, R., Mazein, A., Auffray, C., Babur, Ö., et al. (2021). Newt: a comprehensive web-based tool for viewing, constructing and analyzing biological maps. Bioinformatics 37:1475–1477.

Berardini, T. Z., Reiser, L., Li, D., Mezheritsky, Y., Muller, R., Strait, E., and Huala, E. (2015). The arabidopsis information resource: Making and mining the “gold standard” annotated reference plant genome. genesis 53:474–485.

Bergmann, F. T., Czauderna, T., Dogrusoz, U., Rougny, A., Dräger, A., Touré, V., Mazein, A., Blinov, M. L., and Luna, A. (2020). Systems biology graphical notation markup language (SBGNML) version 0.3. J. Integr. Bioinforma. 17.

Bittner, A., Cieśla, A., Gruden, K., Lukan, T., Mahmud, S., Teige, M., Vothknecht, U. C., and Wurzinger, B. (2022). Organelles and phytohormones: a network of interactions in plant stress responses. J. Exp. Bot. 73:7165–7181.

Bornstein, B. J., Keating, S. M., Jouraku, A., and Hucka, M. (2008). LibSBML: an API Library for SBML. Bioinformatics 24:880–881.

Caspi, R., Billington, R., Ferrer, L., Foerster, H., Fulcher, C. A., Keseler, I. M., Kothari, A., Krummenacker, M., Latendresse, M., Mueller, L. A., et al. (2016). The MetaCyc database of metabolic pathways and enzymes and the BioCyc collection of pathway/genome databases. Nucleic Acids Res. 44:D471–D480.

Cheng, C.-Y., Krishnakumar, V., Chan, A. P., Thibaud-Nissen, F., Schobel, S., and Town, C. D. (2017). Araport11: a complete reannotation of the Arabidopsis thaliana reference genome. Plant J. 89:789–804.

Chow, C.-N., Lee, T.-Y., Hung, Y.-C., Li, G.-Z., Tseng, K.-C., Liu, Y.-H., Kuo, P.-L., Zheng, H.-Q., and Chang, W.-C. (2019). PlantPAN3.0: a new and updated resource for reconstructing transcriptional regulatory networks from ChIP-seq experiments in plants. Nucleic Acids Res. 47:D1155–D1163.

Cline, M. S., Smoot, M., Cerami, E., Kuchinsky, A., Landys, N., Workman, C., Christmas, R., Avila-Campilo, I., Creech, M., Gross, B., et al. (2007). Integration of biological networks and gene expression data using Cytoscape. Nat. Protoc. 2:2366–2382.

Cusack, S. A., Wang, P., Lotreck, S. G., Moore, B. M., Meng, F., Conner, J. K., Krysan, P. J., Lehti-Shiu, M. D., and Shiu, S.-H. (2021). Predictive Models of Genetic Redundancy in Arabidopsis thaliana. Mol. Biol. Evol. 38:3397–3414.

Eckardt, N. A. (2015). The Plant Cell Reviews Dynamic Aspects of Plant Hormone Signaling and Crosstalk. Plant Cell 27:1–2.

Edmonds, J., and Karp, R. M. (1972). Theoretical Improvements in Algorithmic Efficiency for Network Flow Problems. J. ACM 19:248–264.

Foix, L., Nadal, A., Zagorščak, M., Ramšak, Ž., Esteve-Codina, A., Gruden, K., and Pla, M. (2021). Prunus persica plant endogenous peptides PpPep1 and PpPep2 cause PTI-like transcriptome reprogramming in peach and enhance resistance to Xanthomonas arboricola pv. pruni. BMC Genomics 22:360.

Gansner, E. R., and North, S. C. (2000). An open graph visualization system and its applications to software engineering. Softw. Pract. Exp. 30:1203–1233.

Garrett, K. A. (2013). Agricultural impacts: Big data insights into pest spread. Nat. Clim. Change 3:955–957.

Hagberg, A. A., Schult, D. A., and Swart, P. J. (2008). Exploring Network Structure, Dynamics, and Function using NetworkX. In Proceedings of the 7th Python in Science Conference (SciPy 2008), p. Pasadena, CA.

Hassani-Pak, K., Singh, A., Brandizi, M., Hearnshaw, J., Parsons, J. D., Amberkar, S., Phillips, A. L., Doonan, J. H., and Rawlings, C. (2021). KnetMiner: a comprehensive approach for supporting evidence-based gene discovery and complex trait analysis across species. Plant Biotechnol. J. 19:1670–1678.

Hastings, J., Owen, G., Dekker, A., Ennis, M., Kale, N., Muthukrishnan, V., Turner, S., Swainston, N., Mendes, P., and Steinbeck, C. (2016). ChEBI in 2016: Improved services and an expanding collection of metabolites. Nucleic Acids Res. 44:D1214–D1219.

Herwig, R., Hardt, C., Lienhard, M., and Kamburov, A. (2016). Analyzing and interpreting genome data at the network level with ConsensusPathDB. Nat. Protoc. 11:1889–1907.

Hunter, M. C., Smith, R. G., Schipanski, M. E., Atwood, L. W., and Mortensen, D. A. (2017). Agriculture in 2050: Recalibrating targets for sustainable intensification. BioScience 67:386–391.

IPPC Secretariat (2021). Scientific review of the impact of climate change on plant pests – A global challenge to prevent and mitigate plant pest risks in agriculture, forestry and ecosystems. Rome: FAO on behalf of the IPPC Secretariat.

Juteršek, M., Petek, M., Ramšak, Ž., Moreno-Giménez, E., Gianoglio, S., Mateos-Fernández, R., Orzáez, D., Gruden, K., and Baebler, Š. (2022). Transcriptional deregulation of stress-growth balance in *Nicotiana benthamiana* biofactories producing insect sex pheromones. Front. Plant Sci. 13.

Kanehisa, M., Sato, Y., Kawashima, M., Furumichi, M., and Tanabe, M. (2016). KEGG as a reference resource for gene and protein annotation. Nucleic Acids Res. 44:D457–D462.

Keating, S. M., Waltemath, D., König, M., Zhang, F., Dräger, A., Chaouiya, C., Bergmann, F. T., Finney, A., Gillespie, C. S., Helikar, T., et al. (2020). SBML Level 3: an extensible format for the exchange and reuse of biological models. Mol. Syst. Biol. 16:e9110.

Keiichiro Ono, Jorge Bouças, Kozo Nishida, and Barry Demchak Py4cytoscape. https://py4cytoscape.readthedocs.io/

Klarner, H., Streck, A., and Siebert, H. (2017). PyBoolNet: A python package for the generation, analysis and visualization of boolean networks. Bioinformatics 33:770–772.

Knight, M. R., Smith, S. M., and Trewavas, A. J. (1992). Wind-induced plant motion immediately increases cytosolic calcium. Proc. Natl. Acad. Sci. 89:4967–4971.

König, M. (2020). matthiaskoenig/libsbgn-python: 0.2.2 Advance Access published November 1, 2020, doi:10.5281/zenodo.4171366.

Kudla, J., Batistič, O., and Hashimoto, K. (2010). Calcium Signals: The Lead Currency of Plant Information Processing. Plant Cell 22:541–563.

Lichtenberg, J., Yilmaz, A., Welch, J. D., Kurz, K., Liang, X., Drews, F., Ecker, K., Lee, S. S., Geisler, M., Grotewold, E., et al. (2009). The word landscape of the non-coding segments of the Arabidopsis thaliana genome. BMC Genomics 10:463.

Miljkovic, D., Stare, T., Mozetič, I., Podpečan, V., Petek, M., Witek, K., Dermastia, M., Lavrač, N., and Gruden, K. (2012). Signalling Network Construction for Modelling Plant Defence Response. PLoS ONE 7:e51822.

Mueller, L. A., Zhang, P., and Rhee, S. Y. (2003). AraCyc: A Biochemical Pathway Database for Arabidopsis. Plant Physiol. 132:453–460.

Mur, L. A. J., Kenton, P., Atzorn, R., Miersch, O., and Wasternack, C. (2006). The Outcomes of Concentration-Specific Interactions between Salicylate and Jasmonate Signaling Include Synergy, Antagonism, and Oxidative Stress Leading to Cell Death. Plant Physiol. 140:249–262.

Müssel, C., Hopfensitz, M., and Kestler, H. A. (2010). BoolNet -- an R package for generation, reconstruction and analysis of Boolean networks. Bioinformatics 26:1378–1380.

Nomoto, M., Skelly, M. J., Itaya, T., Mori, T., Suzuki, T., Matsushita, T., Tokizawa, M., Kuwata, K., Mori, H., Yamamoto, Y. Y., et al. (2021). Suppression of MYC transcription activators by the immune cofactor NPR1 fine-tunes plant immune responses. Cell Rep. 37.

Otasek, D., Morris, J. H., Bouças, J., Pico, A. R., and Demchak, B. (2019). Cytoscape Automation: empowering workflow-based network analysis. Genome Biol. 20:185.

Peixoto, T. P. (2014). The graph-tool python library. figshare Advance Access published 2014, doi:10.6084/m9.figshare.1164194.

Pirayesh, N., Giridhar, M., Ben Khedher, A., Vothknecht, U. C., and Chigri, F. (2021). Organellar calcium signaling in plants: An update. Biochim. Biophys. Acta BBA - Mol. Cell Res. 1868:118948.

Podpečan, V. (2023). vpodpecan/pysbgn: v0.2.1 Advance Access published May 24, 2023, doi:10.5281/zenodo.7966410.

Podpečan, V., Ramšak, Ž., Gruden, K., Toivonen, H., and Lavrač, N. (2019). Interactive exploration of heterogeneous biological networks with Biomine Explorer. Bioinformatics 35:5385–5388.

Pylianidis, C., Osinga, S., and Athanasiadis, I. N. (2021). Introducing digital twins to agriculture. Comput. Electron. Agric. 184:105942.

Ramšak, Ž., Baebler, Š., Rotter, A., Korbar, M., Mozetič, I., Usadel, B., and Gruden, K. (2014). GoMapMan: integration, consolidation and visualization of plant gene annotations within the MapMan ontology. Nucleic Acids Res. 42:D1167–D1175.

Ramšak, Ž., Coll, A., Stare, T., Tzfadia, O., Baebler, Š., Van de Peer, Y., and Gruden, K. (2018). Network modeling unravels mechanisms of crosstalk between ethylene and salicylate signaling in potato. Plant Physiol. 178:488–499.

Rentel, M. C., and Knight, M. R. (2004). Oxidative Stress-Induced Calcium Signaling in Arabidopsis. Plant Physiol. 135:1471–1479.

Rocha-Sosa, M., Sonnewald, U., Frommer, W., Stratmann, M., Schell, J., and Willmitzer, L. (1989). Both developmental and metabolic signals activate the promoter of a class I patatin gene. EMBO J. 8:23–29.

Sebastian Kalinowski, Peter Nowee, and Ero Carrera (2023). pydot Advance Access published May 16, 2023.

Shannon, P., Markiel, A., Ozier, O., Baliga, N. S., Wang, J. T., Ramage, D., Amin, N., Schwikowski, B., and Ideker, T. (2003). Cytoscape: A Software Environment for Integrated Models of Biomolecular Interaction Networks. Genome Res. 13:2498–2504.

Shukla, P. R., Skea, J., Slade, R., Al Khourdajie, A., van Diemen, R., McCollum, D., Pathak, M., Some, S., Vyas, R., Fradera, M., et al. eds. IPCC, 2022: Summary for Policymakers. In Climate Change 2022: Mitigation of Climate Change. Contribution of Working Group III to the Sixth Assessment Report of the Intergovernmental Panel on Climate Change, p. Cambridge, UK and New York, NY, USA: Cambridge University Press.

Steinwand, M. A., and Ronald, P. C. (2020). Crop biotechnology and the future of food. Nat. Food 1:273–283.

Szklarczyk, D., Kirsch, R., Koutrouli, M., Nastou, K., Mehryary, F., Hachilif, R., Gable, A. L., Fang, T., Doncheva, N. T., Pyysalo, S., et al. (2023). The STRING database in 2023: protein–protein association networks and functional enrichment analyses for any sequenced genome of interest. Nucleic Acids Res. 51:D638–D646.

Wilkinson, M. D., Dumontier, M., Aalbersberg, Ij. J., Appleton, G., Axton, M., Baak, A., Blomberg, N., Boiten, J.-W., da Silva Santos, L. B., Bourne, P. E., et al. (2016). The FAIR Guiding Principles for scientific data management and stewardship. Sci Data 3:160018.

Tracy, F. E., Gilliham, M., Dodd, A. N., Webb, A. a. R., and Tester, M. (2008). NaCl-induced changes in cytosolic free Ca2+ in Arabidopsis thaliana are heterogeneous and modified by external ionic composition. Plant Cell Environ. 31:1063–1073.

Tyanova, S., Temu, T., Sinitcyn, P., Carlson, A., Hein, M. Y., Geiger, T., Mann, M., and Cox, J. (2016a). The Perseus computational platform for comprehensive analysis of (prote)omics data. Nat. Methods 13:731–740.

Tyanova, S., Temu, T., and Cox, J. (2016b). The MaxQuant computational platform for mass spectrometry-based shotgun proteomics. Nat. Protoc. 11:2301–2319.

United Nations Department of Economic and Social Affairs (UN DESA), Population Division (2022). World Population Prospects 2022: Summary of Results.

Van Bel, M., Silvestri, F., Weitz, E. M., Kreft, L., Botzki, A., Coppens, F., and Vandepoele, K. (2022). PLAZA 5.0: extending the scope and power of comparative and functional genomics in plants. Nucleic Acids Res. 50:D1468–D1474.

Zagorščak, M., Blejec, A., Ramšak, Ž., Petek, M., Stare, T., and Gruden, K. (2018). DiNAR: revealing hidden patterns of plant signalling dynamics using Differential Network Analysis in R. Plant Methods 14:78.

Zhang, N., Zhou, S., Yang, D., and Fan, Z. (2020). Revealing Shared and Distinct Genes Responding to JA and SA Signaling in Arabidopsis by Meta-Analysis. Front. Plant Sci. 11.

