## Supplementary material for "Stress Knowledge Map: A knowledge graph resource for systems biology analysis of plant stress responses": S03_SupplementaryFigure1_Motifs-AtRD29A-StRD29.pdf

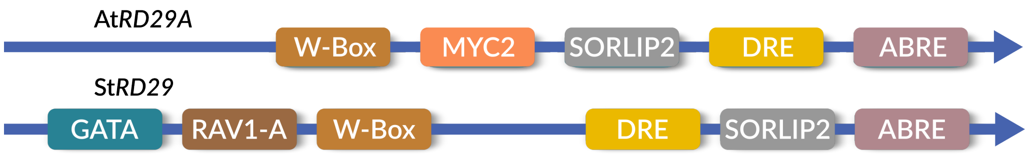


**Supplementary Figure 1: Visualisation of abiotic stress related cis-regulatory binding motifs within the 1 kbp upstream region of the transcription initiation site of AtRD29A and StRD29.** ABRE: ABA-Responsive Element; DRE: Dehydration Responsive Elementi; GATA-Box: light responsive GATA factor–binding motif; SORLIP2: Sequence Over-Represented in Light-Induced Promoters; RAV1-A: RAV1 binding sequence; W-Box: WRKY recognition element; MYC2: basic-helix-loop-helix transcription factor MYC2 binding site.
