## Supplementary material for "Stress Knowledge Map: A knowledge graph resource for systems biology analysis of plant stress responses": S04_SupplementaryFigure2_ABA-response-of-StRD29.pdf

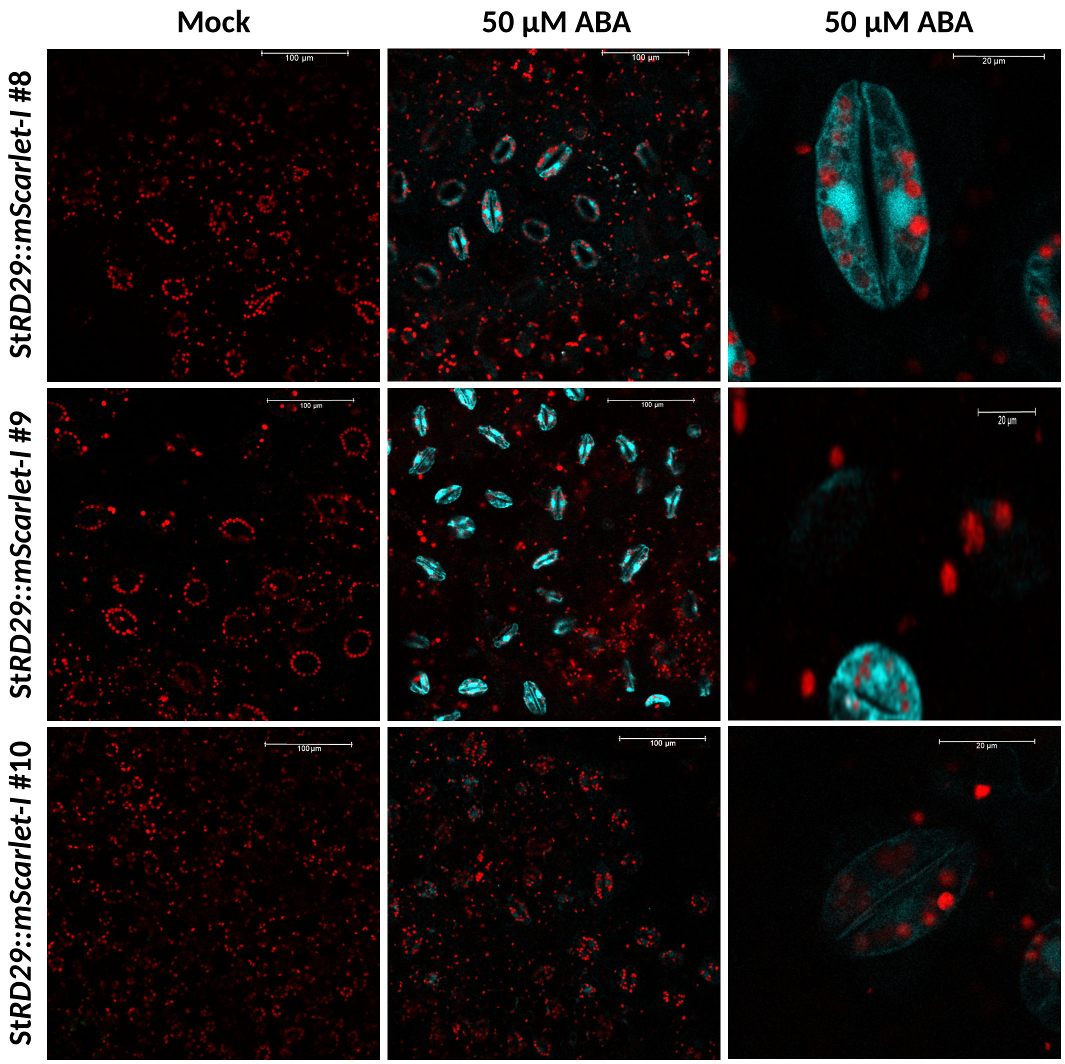


**Supplementary Figure 2: Microscopic analyses of the ABA response of St*RD29::mScarlet-I*.** The fluorophore mScarlet-I was expressed under the control of the StRD29 promoter in transgenic potato plants. Leaf discs (7 mm) of three different St*RD29::mScarlet-I* lines (#10 with very weak response) were treated with 50 µM ABA (or imaging buffer as a mock control) for 24 hours. After the incubation, mScarlet-I fluorescence was visualized with an excitation at 569 nm and emission was recorded at 585-595 nm (cyan) using a Leica SP8 lightning. Chlorophyll fluorescence (red) was visualized with an excitation at 569 nm with emission recorded at 650 – 705 nm.
