## Supplementary material for "Stress Knowledge Map: A knowledge graph resource for systems biology analysis of plant stress responses": S11_SupplementaryFigure3_Vector-StRD29-fluc-and-StRD29-mScarletI.pdf

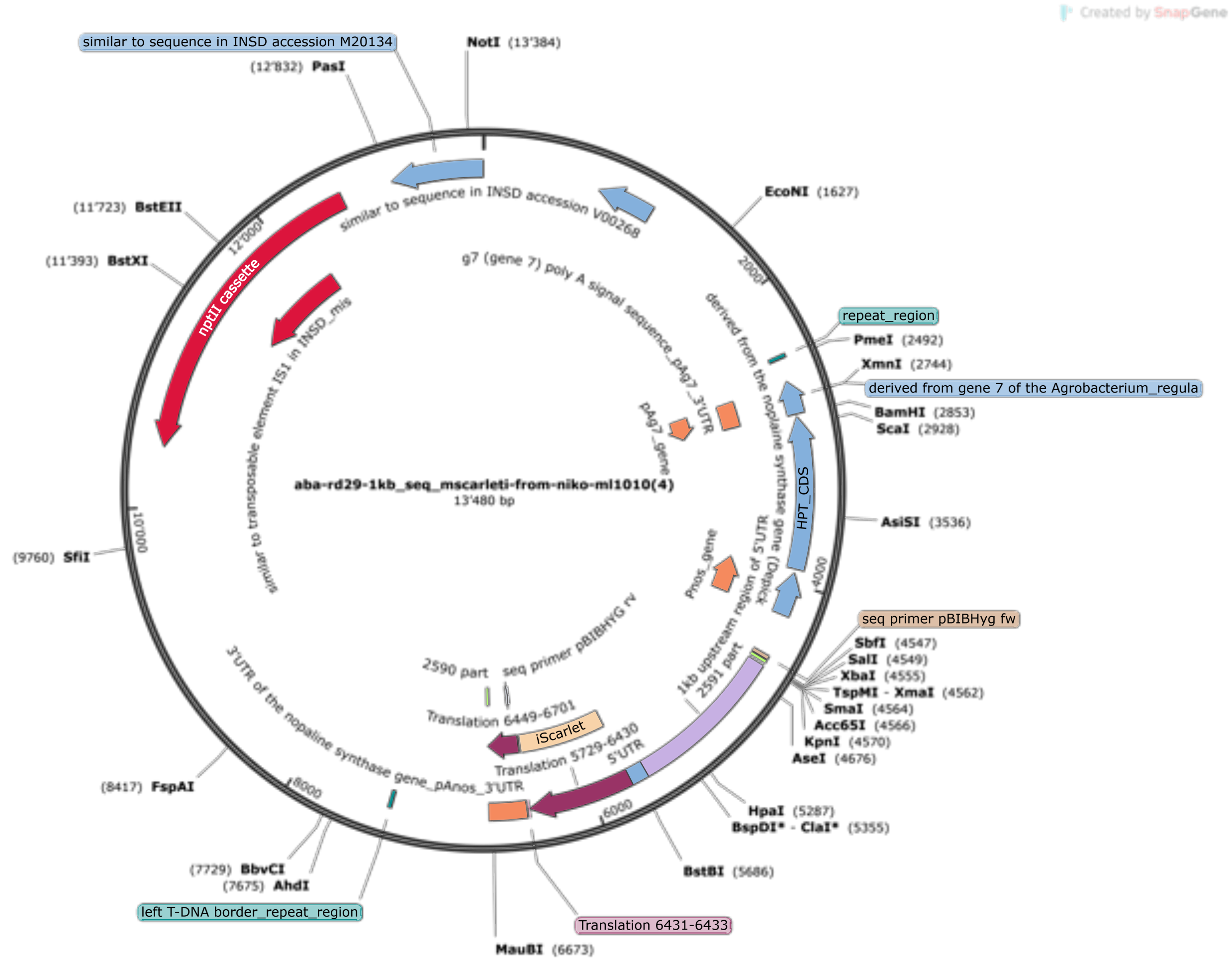


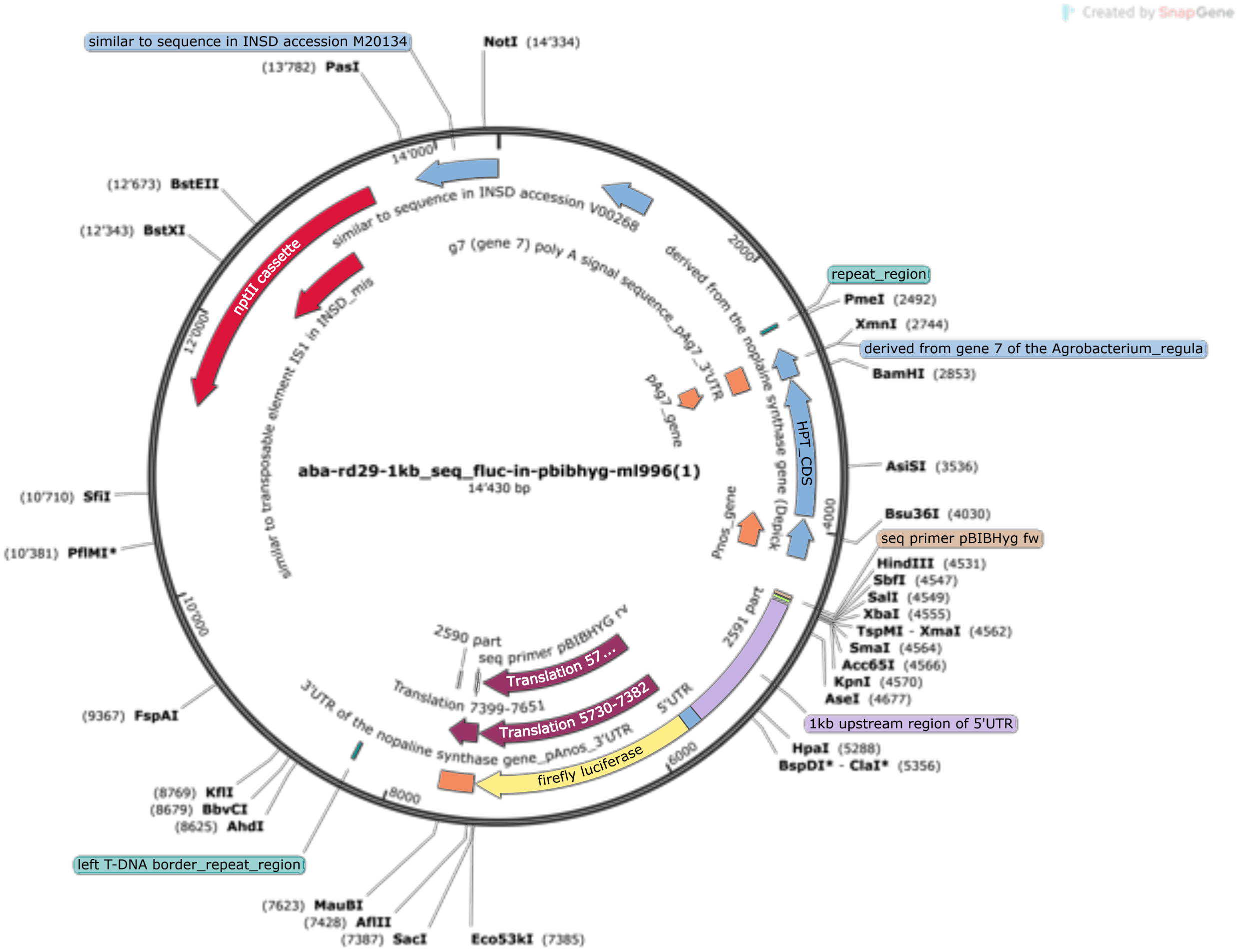


**Supplementary Figure 3:** Visualisation of the features of Vector pBibHyg carrying StRD29::mScarletI. (above) and StRD29::fluc (below).
